# Drug repurposing for ageing research using model organisms

**DOI:** 10.1101/095380

**Authors:** Matthias Ziehm, Satwant Kaur, Dobril K. Ivanov, Pedro J. Ballester, David Marcus, Linda Partridge, Janet M. Thornton

## Abstract

Many increasingly prevalent diseases share a common risk factor: age. However, little is known about pharmaceutical interventions against ageing, despite many genes and pathways shown to be important in the ageing process and numerous studies demonstrating that genetic interventions can lead to a healthier ageing phenotype. An important challenge is to assess the potential to repurpose existing drugs for initial testing on model organisms, where such experiments are possible. To this end, we present a new approach to rank drug-like compounds with known mammalian targets according to their likelihood to modulate ageing in the invertebrates *C. elegans* and *Drosophila.* Our approach combines information on genetic effects on ageing, orthology relationships and sequence conservation, 3D protein structures, drug binding and bioavailability. Overall, we rank 743 different drug-like compounds for their likelihood to modulate ageing. We provide various lines of evidence for the successful enrichment of our ranking for compounds modulating ageing, despite sparse public data suitable for validation. The top ranked compounds are thus prime candidates for *in vivo* testing of their effects on lifespan in *C. elegans* or *Drosophila.* As such, these compounds are promising as research tools and ultimately a step towards identifying drugs for a healthier human ageing.

## Introduction

Age is a major risk factor for many increasingly prevalent diseases. Thus, understanding the process of ageing and finding manipulations leading to a healthier ageing phenotype are highly desirable. Many pathways shown to be important in ageing, e.g. the Insulin/IGF-1 signalling pathway, are also central to other biological processes and diseases, e.g. cancer or diabetes. Research into these diseases is often carried out in mammalian systems or cell lines closely related to humans. In particular, drugs are developed mostly for humans and tested in other mammals, where their target proteins are well characterised. Several of these mammalian targets have orthologues in invertebrates which are known to be involved in ageing.

While model organisms closely related to humans would be ideal from the standpoint of transferring gained knowledge, the length of time required for observing long-term effects and changes in lifespan and other differences in ageing phenotypes are often prohibitive, for both ethical and financial reasons. Cell cultures are not ideal since mechanisms of cellular senescence are probably distinct from organismal ageing. Instead, ageing research is often done in the invertebrate model organisms *Caenorhabditis elegans* and *Drosophila melanogaster* due to their experimentally more amenable lifespans. Thus, we propose a method to transfer knowledge on small-molecule binding from higher organisms, where data on compound binding is available, to lower organisms, which are common model organisms in ageing, to enable direct testing of the compounds' effects on longevity and ageing. This first step is almost opposite to the more common goal of transferring knowledge from lower to higher organisms, e.g. in drug discovery from mice and rats to humans. Positive effects on ageing in invertebrates would suggest drugs with potential positive effects against human ageing, which would warrant further evaluation in mammalian models. Thus, prioritising and testing such compounds in invertebrates could be a first step towards drug-repurposing for ageing. Overall, the aim is thus to create a list of compounds rank ordered by decreasing likelihood of modulating ageing in *C. elegans* and *D. melanogaster*, which incorporates both likely conserved activity as well as targets that are likely to ameliorate ageing.

Here we propose such a ranking procedure based on information on genes and proteins associated with ageing in different organisms, homology and sequence conservation between them, 3D-protein structures, compound activity information and bioavailability predictions. For each ligand we have produced a report card describing the factors which contribute to its score and subsequent ranking.

## Results

An overview of our approach to identify and rank those ligands most likely to affect ageing in *D. melanogaster* and *C. elegans* is shown in Fig 1. We collected genes and proteins implicated in ageing from various sources as well as their orthologues for *C. elegans, D. melanogaster, M. musculus*, *R. norvegicus, H. sapiens* (see Methods for details). This resulted in a total of 13834 UniProt IDs of ageing-associated proteins (2123 in *C. elegans*, 1864 in *D. melanogaster*, 3663 in *M. musculus*, 1589 in *R. norvegicus* and 4595 in *H. sapiens*, see supplement S1 for a lists of these IDs). In order to estimate whether the compound's activity might be conserved between orthologues of known structure and invertebrate target species, we examined the overall protein conservation, and especially the conservation in the binding site. Therefore, we collected information about 3D protein structures and drug-like molecules shown to bind them. This large data acquisition procedure resulted in 1480 3D protein structures with a bound drug-like ligand. The structure-ligand complexes represent 743 different drug-like compounds fulfilling our data requirement criteria binding to 247 different ageing-associated target proteins.

**Figure 1:**
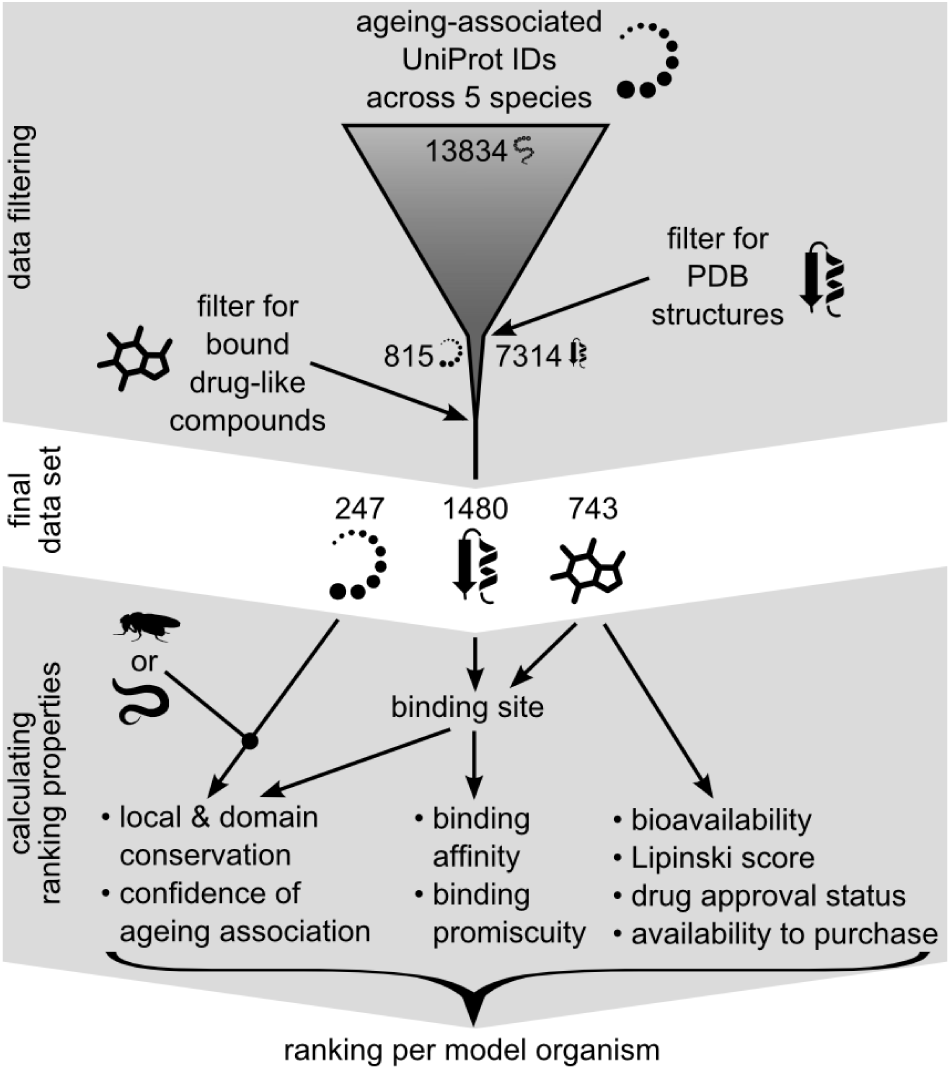
Schematic of principal steps in data gathering and filtering (with number of proteins, structures and drug-like compounds) and the use of these data in determining the components of the ranking procedure. The ranking properties are calculated per model organisms.

Next, we proceeded to rank the ligands using properties of the compounds and their target proteins in the model organism of interest. We developed an empirical scoring function, with the relevant factors multiplicatively linked. In this way, each of the factors is important and cannot be compensated by another factor, as would be the case in an additive scoring function. These factors include relevance to ageing, the conservation of the protein domain or domains containing the binding site, the conservation of the binding site itself, the binding affinity and bioavailability. The resulting base score is then modulated by the following additional terms: drug-likeness, according to Lipinski's rule of five, the promiscuity of the compound, its approval status as a drug, and the availability to purchase the compound. In the methods section we provide the reasoning, technical description and parameter values of each factor and term.

## The ranking

For *D. melanogaster*, 697 compounds, known to bind an ageing-associated protein or an annotated orthologue, were ranked. Compounds scored between 0 (worst) and 1 (best), with a distribution shown in Fig. 2A, with the top 15 compounds scoring above 0.91 and the top 10% scoring above 0.81. For *C. elegans*, 591 compounds were ranked (Fig. 2B), with the top 15 compounds scoring above 0.56 and the top 10% scoring above 0.40.

**Figure 2:**
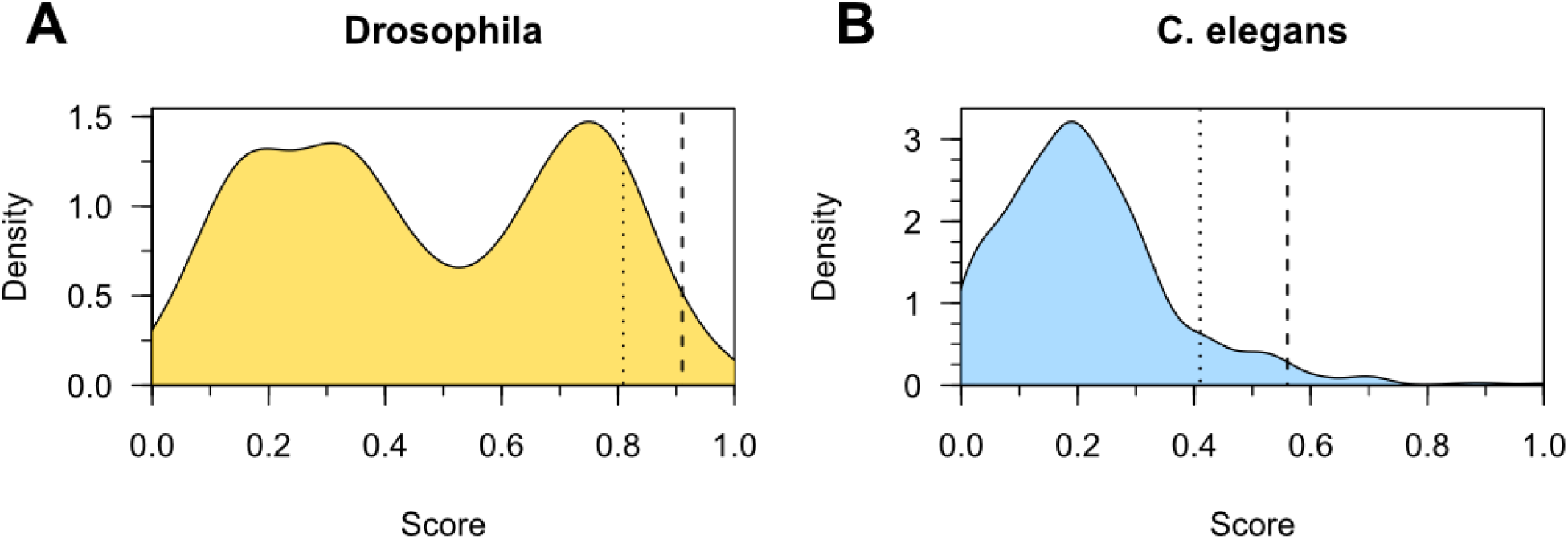
Density distribution of scores in (A) *D. melanogaster* and (B) *C. elegans.* Dashed lines represent the top 15 cut-off and dotted lines the top 10% cut-off.

The overall lower scores for *C. elegans* as compared to *D. melanogaster* originate from two main factors: first, the larger evolutionary distance between *C. elegans* and mammals and, second, the inclusion of the bioavailability predictions score, which is often lower than the arbitrary bioavailability substitute used in *D. melanogaster*, for which such data are not available. The bimodal distribution in *D. melanogaster* is a consequence of the 'ageing implication' score, which has a complex distribution. In *C. elegans* the lower bioavailability values reduce the total scores to give a single peak. Therefore, rankings rather than absolute scores should be compared. Table 1 shows the top 15 ranked compounds for *D. melanogaster* (A) and C. *elegans* (B), of which six (Fig. 3) are ranked highly in both organisms and described briefly below. Several compounds target the same protein and a list of the top 15 compounds targeting different proteins is given in supplement S2. For each compound, we provide a report card including ranking, target protein and its conservation, a graphical representation, images of the 3D compound-target interaction and some additional annotations with links to relevant external resources. The full list of compounds with their respective scores and score components for both *D. melanogaster* and *C. elegans* and all report cards are provided in Supplement S3-S5 and interactively online under https://www.ebi.ac.uk/thornton-srv/software/repurposing/.

**Table 1a:** Top 15 scoring compounds for *D. melanogaster*.

| Rank | PDB<br>HET Code | Name* | Targets** | global identity | binding site identity | Ageing Implication | Domain conservation | Binding site conservation | Binding affinity | Bio-availability | Lipinski Loss | Promiscuity Loss | Purchasability Bonus | Drug approval Bonus | Final Score |
| --- | --- | --- | --- | --- | --- | --- | --- | --- | --- | --- | --- | --- | --- | --- | --- |
| 1 | 1N1 | Dasatinib(Sprycel) | BMX, BTK, MK14 | 67% | 100% | 1. | 0.96 | 1. | 0.95 | 0.9 | 0. | 0. | 0.1 | 0.1 | 1.00 |
| 2 | CT5 | NA | HSP90α | 77% | 92% | 0.98 | 0.97 | 1. | 0.95 | 0.9 | 0. | 0. | 0.1 | 0.08 | 0.99 |
| 3 | I47 | NA | MK14 | 67% | 100% | 1. | 0.96 | 1. | 0.93 | 0.9 | 0. | 0. | 0.1 | 0.08 | 0.98 |
| 4 | SB4 | NA | MK14 | 67% | 100% | 1. | 0.96 | 1. | 0.92 | 0.9 | 0. | 0. | 0.1 | 0.08 | 0.97 |
| 5 | TAK | Dorsomorphin | AMPKα2 | 56% | 100% | 1. | 0.97 | 1. | 0.9 | 0.9 | 0. | 0. | 0.1 | 0.08 | 0.96 |
| 6 | STI | Imatinib(Gleevec) | ABL1, MK14 | 67% | 94% | 1. | 0.97 | 0.99 | 0.88 | 0.9 | 0. | 0. | 0.1 | 0.1 | 0.95 |
| 7 | P01 | Purvalanol | S6Kα1 | 46% | 100% | 1. | 0.96 | 1. | 0.89 | 0.9 | 0. | 0. | 0.1 | 0.08 | 0.95 |
| 8 | P37 | NA | MK14 | 67% | 94% | 1. | 0.96 | 0.95 | 0.94 | 0.9 | 0. | 0. | 0.1 | 0.08 | 0.95 |
| 9 | JNF | NA | MK10 | 62% | 100% | 1. | 0.97 | 1. | 0.87 | 0.9 | 0. | 0. | 0.1 | 0.08 | 0.94 |
| 10 | BAX | Sorafenib(Nexavar) | MK14 | 68% | 95% | 1. | 0.96 | 0.99 | 0.92 | 0.9 | -0.05 | 0. | 0.1 | 0.1 | 0.94 |
| 11 | JNK | NA | MK10 | 62% | 100% | 1. | 0.97 | 1. | 0.92 | 0.9 | -0.05 | 0. | 0.1 | 0.08 | 0.93 |
| 12 | I45 | NA | MK14 | 67% | 88% | 1. | 0.96 | 0.93 | 0.94 | 0.9 | 0. | 0. | 0.1 | 0.08 | 0.93 |
| 13 | BSM | NA | HSP90α | 77% | 88% | 0.98 | 0.97 | 0.93 | 0.94 | 0.9 | 0. | 0. | 0.1 | 0.08 | 0.92 |
| 14 | GVP | NA | AKT2 | 49% | 78% | 1. | 0.96 | 0.91 | 0.93 | 0.9 | 0. | 0. | 0.1 | 0.08 | 0.91 |
| 15 | NIL | Nilotinib(Tasigna) | ABL1, MK11 | 61% | 100% | 1. | 0.97 | 1. | 0.93 | 0.9 | -0.1 | 0. | 0.1 | 0.1 | 0.91 |

**Table 1b:**
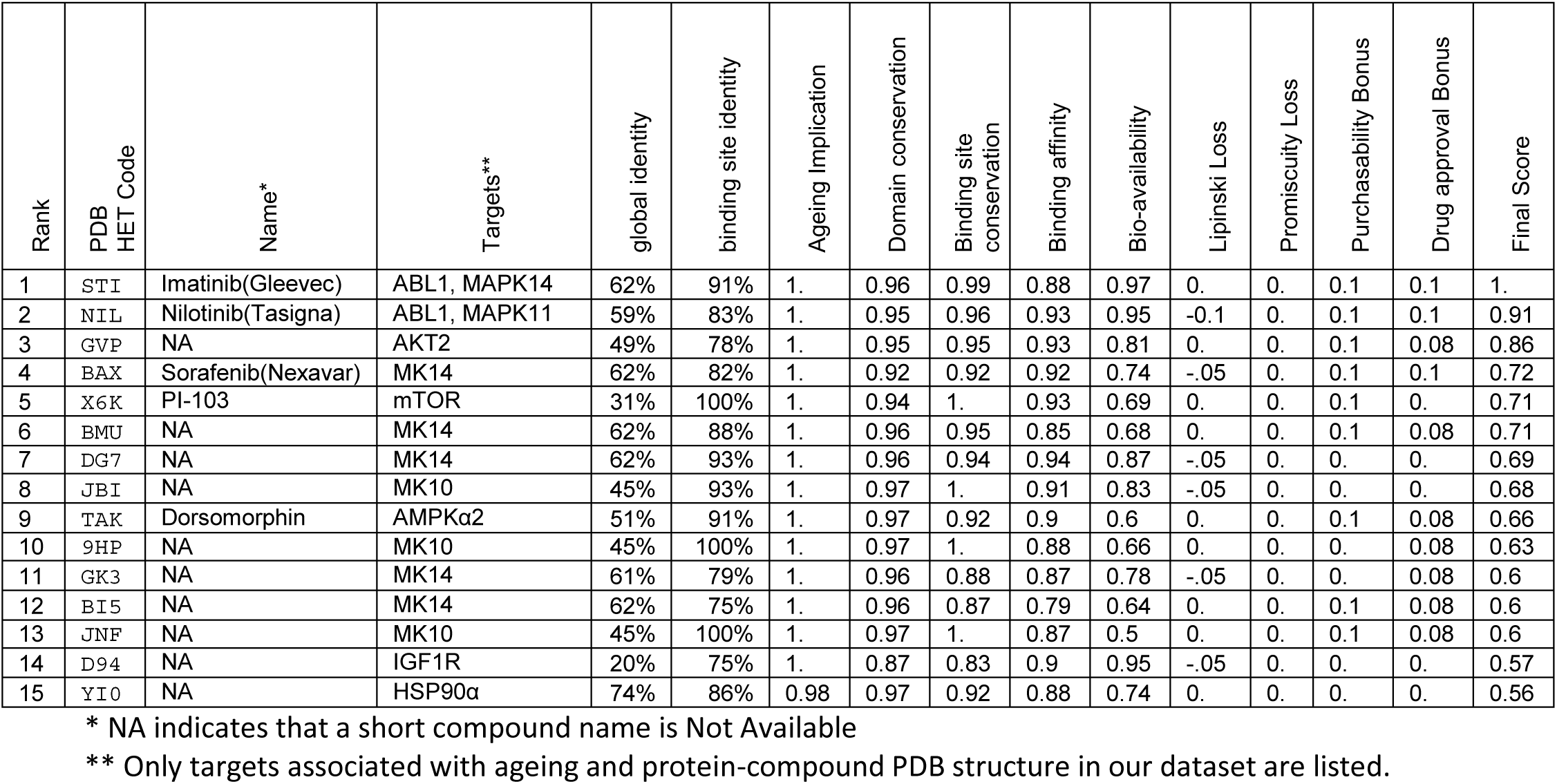
Top 15 scoring compounds for *C. elegans*.

**Figure 3:**
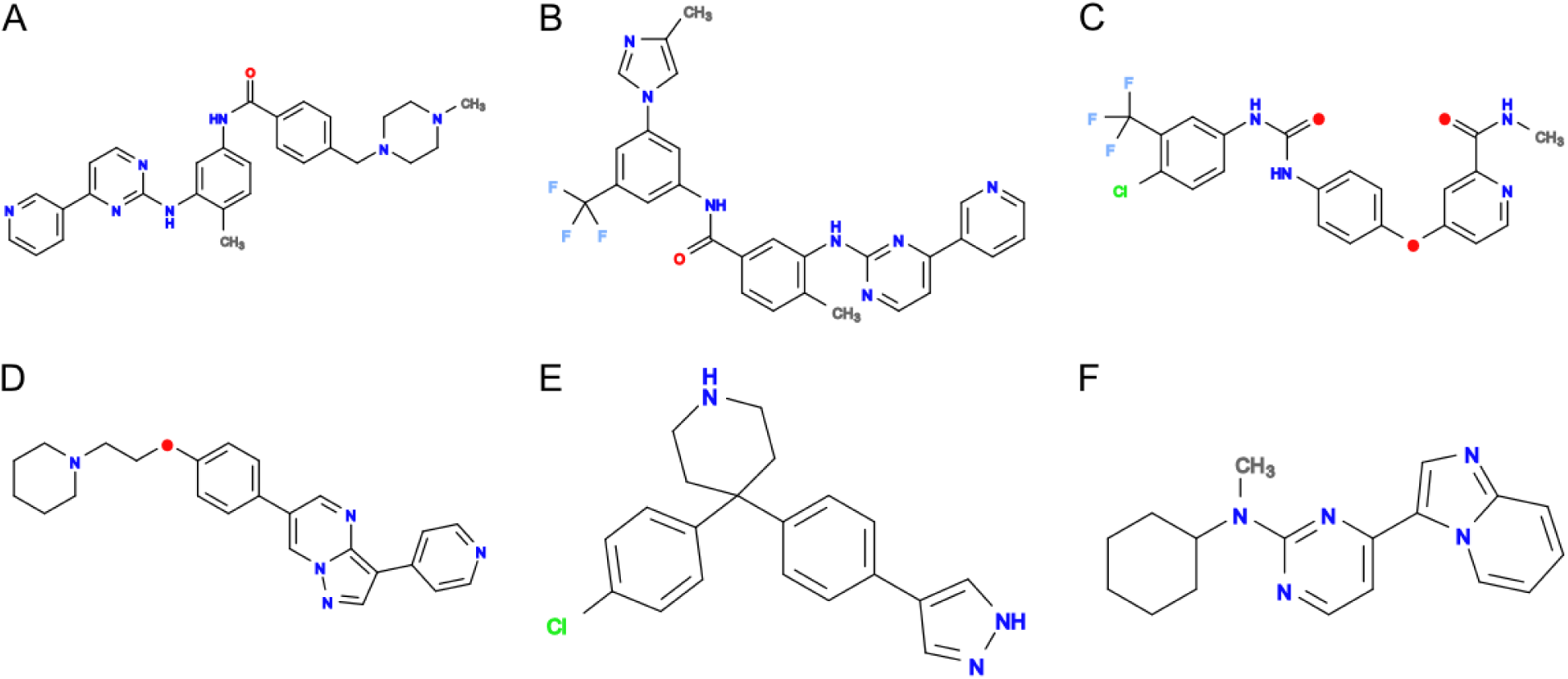
Molecular structures of top overlapping chemical compounds. (A) STI or Imatinib (B) NIL or Nilotinib (C) BAX or Sorafenib (D) TAK or Dorsomorphin/compound c (E) GVP (F) JNF.

The six compounds ranked highly in both organisms exemplify some of the most promising compounds highlighted by the ranking procedure. All six compounds are protein kinase inhibitors, many of them orally available, approved drugs. Others are drug-like probes, not yet approved for the clinic. Despite their common biochemical class, these compounds target a range of different proteins associated with ageing. In general, kinases are often highly conserved between even diverse organisms especially in the compound binding site, which the table shows are even better conserved than the rest of the protein. These compounds rank so highly in part because of the extensive structural work performed for kinases, which is essential for this ranking. In contrast structural data for the membrane bound receptors (which are also often associated with ageing) is scarce and therefore such targets are rarely identified here.

STI, also known as Imatinib or Gleevec (Fig. 3A), is an orally available approved drug for treatment of multiple cancers, especially Philadelphia chromosome-positive chronic myelogenous leukaemia (CML). It is a tyrosine kinase inhibitor targeting a broad range of kinases, amongst which ABL1 is a primary target (Fig. 4A). Human ABL1 has been annotated to be involved in ageing in GenAge release 17 (Tacutu *et al.* 2013). Since kinases all belong to one family, their inhibitors often bind more than one kinase. For example, a second target of Imatinib annotated to be involved in ageing is Mitogen-activated protein kinase 14 (MK14)(Fig. 4B), also known as Mpk2 or p38a in *D. melanogaster*, which is annotated in UniProt with the GO term for determination of adult lifespan (GO:008340). Additionally, there are alleles of the *C. elegans* orthologue pmk-1, which are annotated as lifespan variants in WormBase (Harris *et al.* 2014). The Imatinib binding sites on ABL1 and on MK14 are very well conserved between human and the invertebrates, suggesting a good chance of conserved binding. Additionally, a predicted maximal binding affinity of 100 nM, in mammals is relatively strong and for *C. elegans* the bioavailability prediction indicates very likely successful bioaccumulation in the worm.

**Figure 4:**
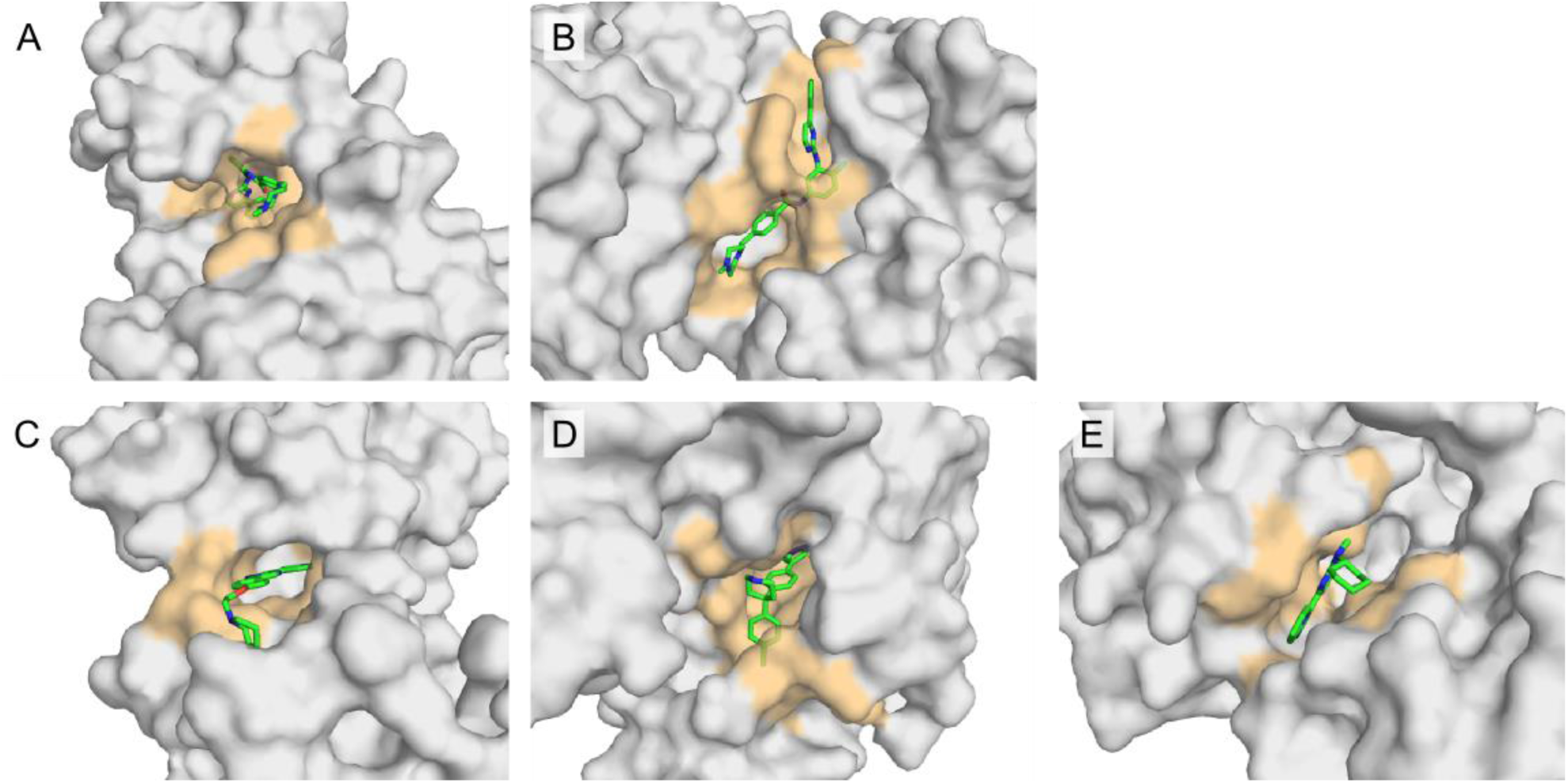
Binding sites of top overlapping compounds. (A) STI or Imatinib binding to human tyrosine kinase ABL1 (B) STI or Imatinib binding to human Mitogen-activated protein kinase 14 (MK14).(C) TAK or Dorsomorphin binding to AMP-activated protein kinase catalytic subunit alpha-2 (D) GVP binding to RAC-beta serine/threonine-protein kinase AKT2 (E) JNF binding to Mitogen-activated protein kinase 10, also known as JNK3.

NIL, also known as Nilotinib or Nexavar (Fig. 3B), is an orally available approved drug for treatment of Imatinib-resistant CML. Like Imatinib, it is an inhibitor targeting a broad range of kinases including ABL1. Nilotinib also binds to Mitogen-activated protein kinase 11 (MK11), while Imatinib binds to MK14. Since MK11 and MK14 are closely related, MK11's role in ageing is implied through the UniProt annotation of the common *D. melanogaster* orthologue Mpk2. Nilotinib has a stronger predicted binding affinity (23 nM) than Imatinib and a similarly good predicted bioavailability in *C. elegans.* However, Nilotinib violates two of the Lipinski Rules, resulting in a slightly lower ranking compared to Imatinib.

BAX, or Sorafenib (Fig. 3C), is another orally available, approved drug for treatment of different cancers, including advanced renal cell carcinoma. Sorafenib is a tyrosine kinase inhibitor of several targets including the Raf kinases and MK14. While the binding site of Sorafenib on MK14 is partly overlapping with that of Imatinib, it makes contact with four more amino acids, and has a similarly strong predicted binding affinity (35 nM). However, this is contrasted by less favourable, albeit still reasonable, predicted bioavailability in C. *elegans* and one violated Lipinski rule.

TAK, also known as Dorsomorphin (Fig. 3D) or compound C, is not an approved drug, but an experimental compound (as listed in DrugBank) and available from a number of vendors. Dorsomorphin has been shown to inhibit Bone morphogenetic protein (BMP) signalling, causing cancer initiating cells to lose some stemcell-like features and induce a proliferation like process (Garulli *et al.* 2014). Dorsomorphin has also been shown to inhibit AMP-activated protein kinase catalytic subunit alpha-2 (Handa *et al.* 2011)(Fig. 4C), which is known as AMPKα or SNF1A in *D. melanogaster* and as aak-2 in *C. elegans.* AMPK is a well-known intracellular energy sensor, involved in the target-of-rapamycin (TOR) signalling pathway. It has also been shown to be involved in ageing in a multitude of experiments in different organism. The binding site is completely conserved between human and *D. melanogaster* and contains only one changed amino acid (Y2H) in *C. elegans.* The predicted binding affinity (63 nM) is quite strong, while the predicted bioavailability for *C. elegans* is only moderate, and no additional information about oral availability is present.

GVP (Fig. 3E) is classified by DrugBank as an experimental antineoplastic agent and commercially available. It inhibits RAC-beta serine/threonine-protein kinase, also known as AKT2 or Protein kinase B beta (Fig. 4D), which is a part of the Insulin/IGF-1 and TOR signalling pathways. The *C. elegans* orthologue akt-2 is annotated in UniProt with the term determination of adult lifespan (GO:008340). More specifically, RNAi inhibition of Akt in *D. melanogaster* and akt-1 and akt-2 in *C. elegans* extends lifespan (Tullet *et al.* 2008; Biteau *et al.* 2010). The binding site contains 4-5 conservative changes (91% similarity between human and either invertebrate). The predicted binding affinity (23nM) is quite strong, with reasonable predicted bioavailability for *C. elegans.*

Finally, JNF (Fig. 3F), is also an experimental compound listed in DrugBank and commercially available. It is a kinase inhibitor targeting the Mitogen-activated protein kinase 10 (MK10) (Fig. 4E), whose *C. elegans* orthologue jnk-1 is annotated in UniProt with the GO term ‘determination of adult lifespan’. More specifically, moderate RNAi of the *D. melanogaster* orthologue Bsk in adult animals extended lifespan, while mutants of *C. elegans* jnk-1 exhibited decreased lifespan (Oh *et al.* 2005; Biteau *et al.* 2010). The reasons for the different lifespan effects, besides possible organismal differences, are likely to be a detrimental developmental effect in *C. elegans* jnk-1 mutants as well as a likely dosage dependency of the effect of modulating MK10. The predicted binding affinity is relatively strong (120 nM), however the predicted bioavailability in *C. elegans* is significantly lower than for the other compounds described above. Overall, these six compounds exemplify the promising candidates for modulating lifespan in both *D. melanogaster* and *C. elegans* highlighted by the ranking procedure.

## Evaluating the ranking

### Modelling & Docking

In the above high-throughput approach, binding of the compounds to the model organism proteins is estimated indirectly, through conservation of sequence. Here we tested this assumption further by modelling the orthologues in *D. melanogaster* and *C. elegans* for one hit (compound P37 to human MK14) and docking the compound directly. P37 is a compound classified by DrugBank as experimental, and does not violate any of the Lipinski rules. Its structure was determined by X-ray crystallography in complex with human MK14 at 2.10Å (PDB:3GFE, Wurz *et al.* 2009). The closest homologues in *D. melanogaster* and C. *elegans* include p38a and pmk-1, respectively, which are 67% and 61% identical. We successfully modelled the protein structures of the homologues having removed the P37 ligand before modelling, not to bias the modelling procedure. P37 was docked into the two models and their binding affinity was predicted 7.65 log Kd/Ki *(D. melanogaster)*, 7.90 log Kd/Ki (C. *elegans).* This affinity was very similar to that predicted from the structure of the human complex: 7.75 log Kd/Ki (Human). This is also evident from a superposition of the binding sites of human crystal structure and *D. melanogaster* or *C. elegans* protein structure model (Fig. 5). The structurally determined conservation in binding is in very good agreement with the sequence-based binding site conservation score of 0.951 *(D. melanogaster)* and 0.914 (C. *elegans)* using the 3D information only to determine the amino acids in contact with the ligand.

**Figure 5:**
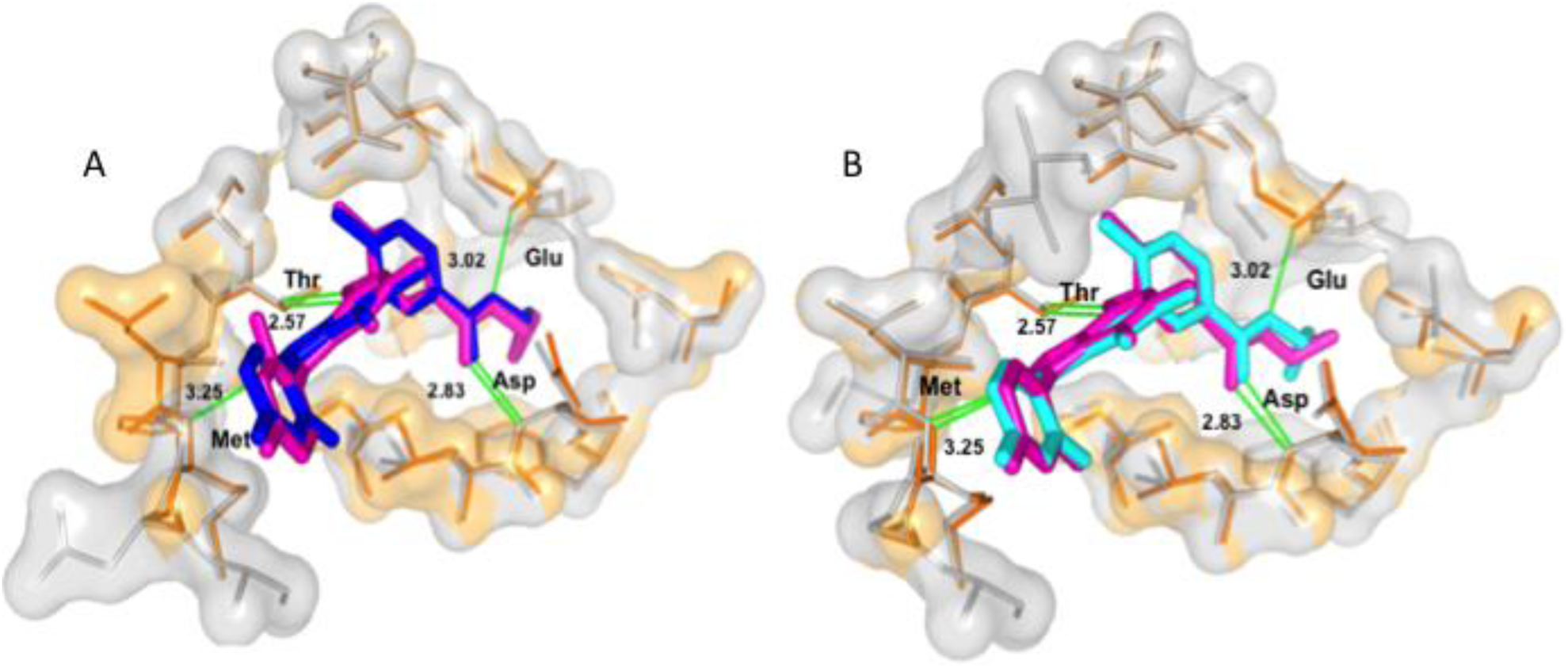
Interaction between the superimposed P37 ligand and the amino acids in the active binding site (A) in human (orange & pink) and *D. melanogaster* (grey & blue) (B) in human (orange & pink) and *C. elegans* (grey and blue).

### Literature mining for the top ranked compounds

We conducted a thorough literature search for articles examining any of the top 25 ranked compounds with respect to lifespan effects. Searching the PubMed database with the compound names and their synonyms in combination with ageing keywords and terms for *C. elegans* and *D. melanogaster* resulted in 38 hits. Querying the PubMed Central database of full text articles resulted in 473 relevant hits of which only four publications describe relevant lifespan experiments, with variable results.

Liu *et al.* (2011) tested Sorafenib (BAX), for effects in Parkinson's models in *C. elegans* and *D. melanogaster.* They reported that Sorafenib at 1 and 10 μM increased survival and reduced locomotor impairment in ddc-GAL4; UAS-G2019S-LRRK2 flies, while none of the concentrations tested was reported to alter lifespan of controls. Furthermore, *C. elegans* treated with Sorafenib showed positive effects on neuronal survival, while overall lifespan was reported unchanged. However, this study tried to exclude effects from cellular processes associated with aging, stating "these effects were maintained to a minimum in our studies" (Liu *et al.* 2011).

Yang *et al.* (2015) evaluated the effect of β-guanidinopropionic acid and Dorsomorphin (TAK) on wild-type D. *melanogaster* lifespan. While the effect of Dorsomorphin on wild type flies was not discussed, the survival curves showed, probably significant, lifespan-shortening, which is in line with what would be expected by inhibiting AMPK and confirms the lifespan changing effects of one of our top ranked compounds.

The third publication tested the lifespan effect of Genistein (4′,5,7-trihydroxyisoflavone) in *C. elegans.* While Genistein is not in our list of compounds, it was found because the highly similar compound 3',4',7-trihydroxyisoflavone (PDB HET code: 47X) is ranked 21^st^ in *C. elegans.* It remains unclear how similar their biological effect is, but Lee *et al.* (2015) report Genistein to significantly increase lifespan and stress tolerance of wild type *C. elegans* lifespan at 50 μM and 100 μM concentrations.

Finally, Wilson *et al.* (2008) reported modest, albeit not significant, beneficial effects (15% increased median lifespan) in *C. elegans* with Piceatannol (PDB HET code: P01, ranked 73^rd^ in *C. elegans).* P01 binds to Ribosomal protein S6 kinase alpha-1, whose binding site is completely conserved between human and invertebrates.

### Rapamycin

Of the well-known drugs examined for their effect on lifespan in invertebrates, only rapamycin (PDB HET code: RAP) is included in our ranking due to the data requirements. Rapamycin was shown to extend lifespan in many organism including *C. elegans*, where it is ranked, however, only 395^th^ according to the method presented here. This is because Rapamycin violates 3 out of the 4 Lipinski rules as well as being predicted by the Burns *et al.* (2010) methodology to have low bioavailability in *C. elegans.* Furthermore, 4 of the 10 amino acids forming the binding site on TOR are not conserved between human and *C. elegans.* However, Rapamycin has a highly unusual mode of action of disrupting a protein-protein interaction between TOR and FKBP and thus also unusual properties. If Rapamycin had a bioavailability score of 0.9, instead of 0.2 it would rank 16^th^ in *C. elegans*, clearly demonstrating the impact of bioavailability. In D. *melanogaster*, with no bioavailability prediction available, Rapamycin ranks 95^th^, classifying it as a promising compound. A different promising kinase inhibitor PI-103 (PDB HET code: X6K), also targeting TOR, is, however, ranked 5^th^ in *C. elegans* and 17^th^ in *D. melanogaster.* In contrast to Rapamycin, PI-103/X6K has no violations of Lipinski's rules for likely drugs, a high predicted bioavailability in *C. elegans*, a completely conserved binding site in both invertebrate species and high predicted binding affinity. While our ranking only considered the effect on TOR, PI-103/X6K also inhibits phosphoinositide 3-kinase (PI3K), which is also strongly associated with ageing (Fan *et al.* 2006), suggesting that it is a promising candidate for trials.

### Unbiased *C. elegans* lifespan screen

Finally, we found a single unbiased screen of a set of compounds for effects on lifespan in *C. elegans* where complete experimental results are reported. The study by Ye *et al.* (2014) tested the Library of Pharmacologically Active Compounds (LOPAC), comprising 1280 different compounds at a fixed concentration of 33μM in liquid culture. Of these compounds only four were included in our *C. elegans* ranking, ranking 220^th^ or lower in our calculations. None of them significantly affected lifespan (p≥0.05) according to Ye *et al.* (2014). Furthermore, 15 of the compounds tested are included in our *D. melanogaster* ranking (the 11 additional compounds had no annotated *C. elegans* orthologues in Compara and thus could not be ranked in *C. elegans).* Three (B43, STR and TDC) of the 15 compounds are reported to significantly affect lifespan (p<0.05), these include the two highest ranked of this list. This is a significant enrichment for lifespan changing compounds by rank (hypergeometric test p<0.03 for overlap of lifespan extending compounds with compounds ranked in the top 250 of the overall list).

## Discussion

In this proof-of-principle study we show that, by combining different types of information, it is possible to create a ranked list of compounds with respect to their likelihood to modulate ageing in invertebrates. The ranking demonstrated strong differences between potential candidates for compound testing compared to random selection of any compound binding an orthologue of a known modulator of ageing.

The pipeline developed is fast and conservative, choosing to focus on those compounds with strongest supporting evidence for an effect on ageing. The pipeline could be 'relaxed' to include more distant orthologues or compounds and proteins without any human complex structural data. Furthermore, the ranking procedure presented here also serves as a blueprint for similar approaches, where individual factors or terms can be omitted or substituted with alternative methods of preference. It is interesting that most of the drugs at the top of the list were developed to target cancers, perhaps reflecting the increased occurrence of cancer with age.

Although no direct experimental validation is available, we provide various lines of evidence indicating the success of the ranking and making the top ranked compounds interesting candidates for experimental *In vivo* testing. Due to uncertainty about compound dosage, such experiments would require testing at multiple concentrations, e.g. three concentrations spanning three orders of magnitude. In *C. elegans*, the use of a specific drug-sensitive mutant strain (partial loss-of-function bus-8 mutant), has been suggested for drug-screening (Partridge *et al.* 2008). A complication in *C. elegans* drug screening can be the live *E. coli* bacteria contained in the most commonly used food, which might metabolise compounds (Cabreiro *et al.* 2013; Zheng *et al.* 2013).

Pharmacological interventions can be an extremely helpful research tool, which enable time-restricted intervention in a dosage dependent way, thereby allowing a more precise control than genetic interventions. Furthermore, various compounds can be easily administered in combination and further combined with established genetic or environmental intervention. As such, pharmacological interventions are an orthogonal manipulation system which will help to further deconvolute the pathways and processes relevant to ageing. The recently established *Caenorhabditis* Intervention Testing Program (CITP) for robustly testing pharmacological interventions for their effect on ageing across different *Caenorhabditis* strains and species further demonstrates the relevance and timeliness of our computational ranking procedure.

Neither *C. elegans* nor *D. melanogaster* are typical model organisms for pharmaceutical research, but successful examples in drug screenings exist (Desalermos *et al.* (2011), Pandey and Nichols (2011)) and demonstrate the feasibility of large-scale compound screens. Clearly, successful evaluation of compounds against ageing in either invertebrate model would require subsequent testing in mammalian models, e.g. in the National Institute on Aging Interventions Testing Program (Warner *et al.* 2000), in order to determine the most likely compounds to influence human ageing.

## Experimental Procedures

### Data sources and mapping

Genes and proteins associated with ageing were obtained from the GenAge database (Tacutu *et al.* 2013) as well as from general databases (UniProt (UniProt Consortium 2014) and Ensembl (Flicek *et al.* 2014)) and organism-specific databases (RGD (Dwinell *et al.* 2009), MGI (Eppig *et al.* 2012), FlyBase (St Pierre *et al.* 2014) and WormBase (Harris *et al.* 2014)) using the Gene Ontology term for ageing (GO:0007568). We considered five selected species of interest: *C. elegans, D. melanogaster, M. musculus, R. norvegicus and H. sapiens.* All results were mapped to the corresponding Ensembl and UniProt entries and all entries were cross-mapped between Ensembl and UniProt. For each gene/protein associated with ageing, all orthologues in the five species were identified using Ensembl Compara. PDB codes corresponding to these proteins were retrieved from UniProt, Ensembl and the PDBsum database (de Beer *et al.* 2014). DrugBank IDs (Law *et al.* 2014) and ChEMBL IDs (Gaulton *et al.* 2012, release 15) of compounds binding ageing-associated proteins were collected from UniProt, PDBsum and ChEMBL. Compound IDs were cross-referenced with UniChem (Chambers *et al.* 2013) and, where possible, mapped to the ZINC (Irwin *et al.* 2012) and eMolecules databases (http://www.emolecules.com/) of commercially available compounds, ChEBI (Hastings *et al.* 2013) and to PDB HET codes, the compound identifiers used in PDB structures. PDB HET codes were used to identify PDB files of ageing-associated proteins with bound drug-like ligands of the DrugBank and ChEMBL set and at least one orthologue in *C. elegans* or *D. melanogaster.* All targets with no homologue in *C. elegans* or *D. melanogaster* were filtered out, because these are the target species. Additionally, for each homologue-family and species, only the homologues with the smallest number of gaps in the binding site and the highest binding-site identity or similarity were kept.

### The ranking score equations and components

The ranking score is a bounded score between zero and one determined by five multiplicative factors and four additional terms for bonuses and losses, defined by the following formula:

Ranking score = max(min(Ageing implication × Domain conservation × Binding site conservation × Binding affinity × Bioavailability + Lipinski loss + Promiscuity loss + Availability to purchase bonus + Approved drug bonus, 1),0).

#### Ageing implication

This factor represents the certainty of our knowledge associating the protein target with ageing. It is derived from the Gene Ontology (Ashburner *et al.* 2000) annotation evidence codes. A distribution of scores for this and the other terms is available online under https://www.ebi.ac.uk/thornton-srv/software/repurposing/.

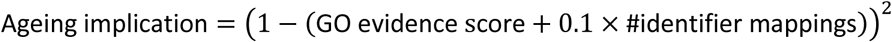

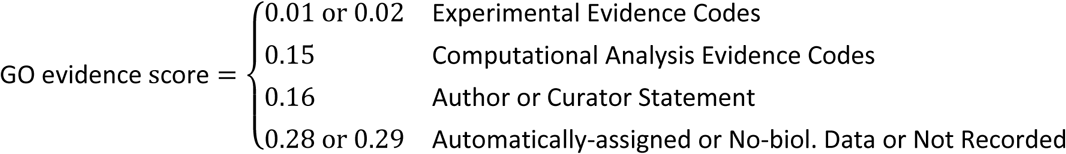

Genes implied through being listed in GenAge are assigned a GO evidence score of 0.01.

#### Protein domain conservation

This factor and the factor for binding site conservation represent the requirements for conservation to maintain binding of the compound to the protein target. In practice, we use a conservation score for the protein domains, which contain the binding site contacts. A multiple sequence alignment (MSA) of the amino acid sequence from the 3D structure, the corresponding UniProt entries, and their homologues in the five species of interest was constructed using MAFFT with maxiterate 1000 and localpair options (Katoh & Toh 2008). Pairwise sequence identity between the protein of known structure and each homologue, as well as the pairwise Grantham-based similarity (Grantham 1974) were calculated. Grantham-based similarity was defined as 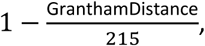 resulting in a 0 to 1 scaled similarity measure. Values for amino acid codes B, Z and X were calculated as average of the D and N, Q and E and all amino acids respectively (see Supplement S6 for Grantham-based similarity matrix). We used protein domain assignments from Gene3D (Lees *et al.* 2014) for the UniProt of the 3D structure. All domains in contact with the bound ligand where jointly considered. For amino acids contacts, which lay outside annotated domains, we used a window of ±50 amino acids around the contact instead.

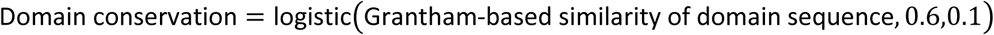

The logistic transformation, 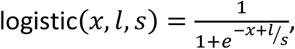 was applied, because differences in medium similarities are more important than equally large differences among very low or very high percentage similarities. For compounds binding to more than one protein, the maximum values for domain conservation, binding-site conservation and binding affinity were used.

#### Binding site conservation

This factor represents the conservation in the binding site, which is especially important for conserved activity of the compound. We obtained the amino acid positions from the MSA that are in contact with the bound ligand as indicated in PDBsum. Since we observed that a large majority of binding sites contained more than 50% identical amino acid residues, we based this factor on the Grantham-based similarity score of 50% most dissimilar positions only, to increase sensitivity of the factor.

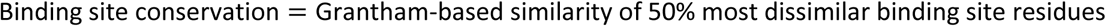

#### Binding affinity

While all compounds ranked are shown to bind their target, as evident from a ligand-protein complex structure, the binding affinity of the interaction can range from low to high. Clearly, a compound binding only with millimolar affinity (10^-3^M) to a target is less likely to bind a slightly altered target than a compound binding with nanomolar affinity (10^-9^M). This factor represents the binding affinity and is derived from the −log binding affinity predicted by RF-Score v2 (Ballester *et al.* 2014) based on the protein-ligand complex structure in 3D. Predicted binding affinities were used since measured binding affinities were not available for all complexes and previous validation demonstrated the high accuracy of RF-Score at this task (Ain *et al.* 2015). Since RF-Score v2 required input files in PDBbind format, but not all complexes of interest were in the PDBbind database (Wang *et al.* 2004), we created and successfully tested a PDBbind format mimicking pipeline (see Supplement S7).

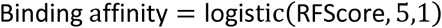

#### Bioavailability

In order for any compound to be able to exert an effect, it needs be able to interact with its, mostly intracellular, targets. Thus, bioavailability is an essential prerequisite. In *C. elegans* bioavailability has been found to be very limited (<10%) in standard approaches (Burns *et al.* 2010). After evaluating the only published bioavailability predictor for *C. elegans* (Burns *et al.* 2010) for its predictive performance (see Supplement S7 for details), we used its score logistically transformed and scaled as a basis for our bioavailability score in *C. elegans.* Since we could not successfully evaluate the relevance or suitability of the predictor for *D. melanogaster*, we opted not to use these predictions in the *D. melanogaster* ranking, but rather substitute the bioavailability score with a fixed term, which can be readily replaced when predictions or measurements of bioavailability in *D. melanogaster* becomes available.

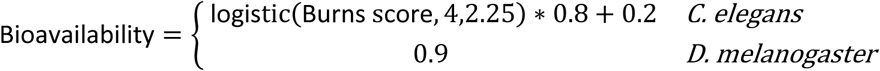

The modulating losses and bonuses are defined as follows:

#### Lipinski score

This term is one of four terms representing non-essential beneficial and detrimental properties. This term has been used to represent the drug likeness based on Lipinski's rule of five (Lipinski *et al.* 2001), which has been widely applied in drug development processes. This rule of thumb states that compounds have a high likelihood of being orally active in humans if they have a molecular weight less than 500 Daltons, an octanol-water partition coefficient log P of no more than 5, no more than 5 Hydrogen-bond donors and no more than 10 Hydrogen-bond acceptors. This and the following terms are scaled to modulate the overall score.

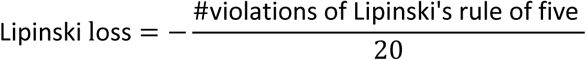

#### Binding promiscuity

An examination of the list of compounds and their targets showed a few compounds, which are found to bind to a large number of targets. These compounds included well-known omnipresent metabolites such as ATP, but also compounds often used to aid crystallisation processes, such as PEG. These highly promiscuous compounds are less interesting compared to more specific compounds targeting one or a few targets. This term penalises compounds by number of targets.

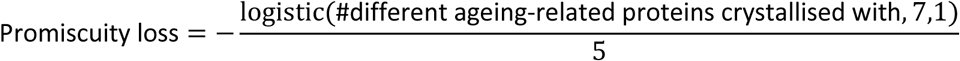

#### Availability to purchase

The final two terms of the ranking are not requirements for the successful modulation of ageing, but influence the ease of performing any experiments and taking positive results forward. This term gives a bonus for purchasable compounds, which is a requirement for most researchers in order to examine its effects, as few will have the means to synthesise any desired compounds.

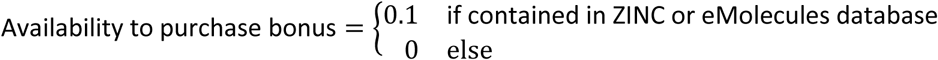

#### Drug approval status

Finally, approved human drugs and drug candidates (as defined in DrugBank (Law *et al.* 2014)), receive a bonus. These compounds would be especially attractive if found to extend lifespan in model organisms since they have already been shown to be tolerated by humans at least under certain circumstances. Thus, they might offer shorter development routes to beneficial interventions against human ageing by drug-repurposing.

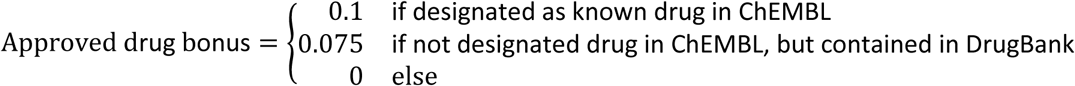

### Protein structure modelling and docking

Single template modelling was performed using Automodel and loop modelling from Modeller 9.13 (Eswar *et al.* 2006). Trailing amino acid residues lacking structural information and the P37 ligand were removed before modelling, while other hetero atoms such as ions were included. The best model was selected based on Discrete Optimized Protein Energy (DOPE) using the CHARM22 force field and Root Mean Square deviation (RMSD). Rigid-protein, flexible-ligand docking experiments were performed using Autodock vina4.2 (Trott & Olson 2010) with a 27.5Å search box. Finally, protein structure model files with docked ligand were converted to PDBbind mimicking files for RF-Score binding affinity prediction.

### Literature mining

Literature mining was performed by querying PubMed and PubMed Central (NCBI Resource Coordinators 2015) using the NCBI Entrez Programming Utilities (eUtils; http://eutils.ncbi.nlm.nih.gov/entrez/eutils). We created search term dictionaries for compound names and synonyms, lifespan terms, and species terms, with all combinations of a drug dictionary term, a survival dictionary term and a species term. The query was conducted to use the automatic term expansion (PubMed), which, for example, expanded “ageing” to “(aging[MeSH Term] OR aging[All Fields] OR ageing[All Fields])”, or manual equivalent expansion (PubMed Central). The compound dictionary was constructed to contain for each of the top 25 compounds for D. *melanogaster* and *C. elegans:* the ChEMBL, DrugBank and ZINC identifiers if available, all names and synonyms listed by these three resources as well as all names from the NCI/CADD Chemical Identifier Resolver (http://cactus.nci.nih.gov/chemical/structure). Names shorter than three characters were excluded. The survival dictionary comprised “longevity”, “lifespan”, “life-span”, “life span”, “life history”, “survival”, “mortality”, “ageing” and the species dictionary included “Drosophila melanogaster”, “Drosophila”, “melanogaster”, “Caenorhabditis”, “elegans”, “C.elegans”. The dictionaries were constructed to be relatively promiscuous in order not to miss any relevant publications. For the PubMed Central queries, the survival dictionary did not contains “ageing”, but all queries had the additional constraint “+AND+(aging[MH]+OR+aging[ARTICLE]+OR+ageing[ARTICLE])” to gear the results more toward longevity experiments in contrast to cancer survival. In order to avoid finding articles where the relevant terms are only part of the references, we restricted the survival and species terms, but not the compound term, to the article body. For all PubMed and PubMed Central queries, all space characters were substituted with '+' as required by the query tool.

## Acknowledgements

We like to thank Roman Laskowski for computational access to PDBsum, Rita Santos for help with calculating the bioavailability predictions, Thomas Holder (Schrödinger Inc) for help with the automated creation of the 3D binding site representations, CHEBI team for curation of highly ranked compounds not previously included. The whole Thornton group and Institute of Healthy Ageing for helpful discussion and advice.

This work was funded by EMBL (M.Z., J.M.T.), Brunel University (S.K.) and the Wellcome Trust [098565/Z/12/Z](M.Z., D.K.I., L.P., J.M.T).

## Author Contributions

M.Z. and J.M.T. designed the study. M.Z. and S.K. collected the data, developed and applied the analyses with the help of D.K.I., P.J.B. and D.M.. M.Z., J.M.T. and L.P. interpreted the results and wrote the manuscript. All authors read, revised and approved the final version of this manuscript.

## Supporting Information Listing

S1: Table of ageing-associated UniProt IDs.

S2: Table of Top 15 scoring compounds for *D. melanogaster* and *C. elegans* targeting at least one protein implied in ageing not targeted by a higher ranked compound.

S3: Table of full *D. melanogaster* ranking including individual scores.

S4: Table of full *C. elegans* ranking including individual scores.

S5: Zip-Archive of all report cards.

S6: Grantham based similarity matrix.

S7: Supplemental Text: PDBbind mimicking files for RF-Score & Bioavailability Prediction assessment.

## References

Ain QU, Aleksandrova A, Roessler FD, Ballester PJ (2015). Machine-learning scoring functions to improve structure-based binding affinity prediction and virtual screening. Wiley interdisciplinary reviews. Computational molecular science. 5, 405–424.

Ashburner M, Ball CA, Blake JA, Botstein D, Butler H, Cherry JM, Davis AP, Dolinski K, Dwight SS, Eppig JT, Harris MA, Hill DP, Issel-Tarver L, Kasarskis A, Lewis S, Matese JC, Richardson JE, Ringwald M, Rubin GM, Sherlock G (2000). Gene ontology: tool for the unification of biology. The Gene Ontology Consortium. Nat Genet. 25, 25–29.

Ballester PJ, Schreyer A, Blundell TL (2014). Does a more precise chemical description of protein-ligand complexes lead to more accurate prediction of binding affinity? J Chem Inf Model. 54, 944–955.

Biteau B, Karpac J, Supoyo S, Degennaro M, Lehmann R, Jasper H (2010). Lifespan extension by preserving proliferative homeostasis in Drosophila. PLoS Genet. 6, e1001159–e1001159.

Burns AR, Wallace IM, Wildenhain J, Tyers M, Giaever G, Bader GD, Nislow C, Cutler SR, Roy PJ (2010). A predictive model for drug bioaccumulation and bioactivity in Caenorhabditis elegans. Nat Chem Biol. 6, 549–557.

Cabreiro F, Au C, Leung KY, Vergara-Irigaray N, Cocheme HM, Noori T, Weinkove D, Schuster E, Greene ND, Gems D (2013). Metformin retards aging in C. elegans by altering microbial folate and methionine metabolism. Cell. 153, 228–239.

Chambers J, Davies M, Gaulton A, Hersey A, Velankar S, Petryszak R, Hastings J, Bellis L, McGlinchey S, Overington JP (2013). UniChem: a unified chemical structure cross-referencing and identifier tracking system. J Cheminform. 5, 3–3.

de Beer TA, Berka K, Thornton JM, Laskowski RA (2014). PDBsum additions. Nucleic Acids Res. 42, D292–296.

Desalermos A, Muhammed M, Glavis-Bloom J, Mylonakis E (2011). Using C. elegans for antimicrobial drug discovery. Expert Opin Drug Discov. 6, 645–652.

Dwinell MR, Worthey EA, Shimoyama M, Bakir-Gungor B, DePons J, Laulederkind S, Lowry T, Nigram R, Petri V, Smith J, Stoddard A, Twigger SN, Jacob HJ, Team RGD (2009). The Rat Genome Database 2009: variation, ontologies and pathways. Nucleic Acids Res. 37, D744–D749.

Eppig JT, Blake JA, Bult CJ, Kadin JA, Richardson JE, Group MGD (2012). The Mouse Genome Database (MGD): comprehensive resource for genetics and genomics of the laboratory mouse. Nucleic Acids Res. 40, D881–D886.

Eswar N, Webb B, Marti-Renom MA, Madhusudhan MS, Eramian D, Shen MY, Pieper U, Sali A (2006). Comparative protein structure modeling using Modeller. Current protocols in bioinformatics / editoral board, Andreas D. Baxevanis … [et al.]. Chapter 5, Unit 5 6.

Fan QW, Knight ZA, Goldenberg DD, Yu W, Mostov KE, Stokoe D, Shokat KM, Weiss WA (2006). A dual PI3 kinase/mTOR inhibitor reveals emergent efficacy in glioma. Cancer Cell. 9, 341–349.

Flicek P, Amode MR, Barrell D, Beal K, Billis K, Brent S, Carvalho-Silva D, Clapham P, Coates G, Fitzgerald S, Gil L, Giron CG, Gordon L, Hourlier T, Hunt S, Johnson N, Juettemann T, Kahari AK, Keenan S, Kulesha E, Martin FJ, Maurel T, McLaren WM, Murphy DN, Nag R, Overduin B, Pignatelli M, Pritchard B, Pritchard E, Riat HS, Ruffier M, Sheppard D, Taylor K, Thormann A, Trevanion SJ, Vullo A, Wilder SP, Wilson M, Zadissa A, Aken BL, Birney E, Cunningham F, Harrow J, Herrero J, Hubbard TJ, Kinsella R, Muffato M, Parker A, Spudich G, Yates A, Zerbino DR, Searle SM (2014). Ensembl 2014. Nucleic Acids Res. 42, D749–755.

Garulli C, Kalogris C, Pietrella L, Bartolacci C, Andreani C, Falconi M, Marchini C, Amici A (2014). Dorsomorphin reverses the mesenchymal phenotype of breast cancer initiating cells by inhibition of bone morphogenetic protein signaling. Cellular signalling. 26, 352–362.

Gaulton A, Bellis LJ, Bento AP, Chambers J, Davies M, Hersey A, Light Y, McGlinchey S, Michalovich D, Al-Lazikani B, Overington JP (2012). ChEMBL: a large-scale bioactivity database for drug discovery. Nucleic Acids Res. 40, D1100–D1107.

Grantham R (1974). Amino acid difference formula to help explain protein evolution. Science. 185, 862–864.

Handa N, Takagi T, Saijo S, Kishishita S, Takaya D, Toyama M, Terada T, Shirouzu M, Suzuki A, Lee S, Yamauchi T, Okada-Iwabu M, Iwabu M, Kadowaki T, Minokoshi Y, Yokoyama S (2011). Structural basis for compound C inhibition of the human AMP-activated protein kinase alpha2 subunit kinase domain. Acta crystallographica. Section D, Biological crystallography. 67, 480–487.

Harris TW, Baran J, Bieri T, Cabunoc A, Chan J, Chen WJ, Davis P, Done J, Grove C, Howe K, Kishore R, Lee R, Li Y, Muller HM, Nakamura C, Ozersky P, Paulini M, Raciti D, Schindelman G, Tuli MA, Van Auken K, Wang D, Wang X, Williams G, Wong JD, Yook K, Schedl T, Hodgkin J, Berriman M, Kersey P, Spieth J, Stein L, Sternberg PW (2014). WormBase 2014: new views of curated biology. Nucleic Acids Res. 42, D789–793.

Hastings J, de Matos P, Dekker A, Ennis M, Harsha B, Kale N, Muthukrishnan V, Owen G, Turner S, Williams M, Steinbeck C (2013). The ChEBI reference database and ontology for biologically relevant chemistry: enhancements for 2013. Nucleic Acids Res. 41, D456–463.

Irwin JJ, Sterling T, Mysinger MM, Bolstad ES, Coleman RG (2012). ZINC: A Free Tool to Discover Chemistry for Biology. J Chem Inf Model. 52, 1757–1768.

Katoh K, Toh H (2008). Recent developments in the MAFFT multiple sequence alignment program. Brief Bioinform. 9, 286–298.

Law V, Knox C, Djoumbou Y, Jewison T, Guo AC, Liu Y, Maciejewski A, Arndt D, Wilson M, Neveu V, Tang A, Gabriel G, Ly C, Adamjee S, Dame ZT, Han B, Zhou Y, Wishart DS (2014). DrugBank 4.0: shedding new light on drug metabolism. Nucleic Acids Res. 42, D1091–1097.

Lee EB, Ahn D, Kim BJ, Lee SY, Seo HW, Cha YS, Jeon H, Eun JS, Cha DS, Kim DK (2015). Genistein from Vigna angularis Extends Lifespan in Caenorhabditis elegans. Biomolecules & therapeutics. 23, 77–83.

Lees JG, Lee D, Studer RA, Dawson NL, Sillitoe I, Das S, Yeats C, Dessailly BH, Rentzsch R, Orengo CA (2014). Gene3D: Multi-domain annotations for protein sequence and comparative genome analysis. Nucleic Acids Res. 42, D240–D245.

Lipinski CA, Lombardo F, Dominy BW, Feeney PJ (2001). Experimental and computational approaches to estimate solubility and permeability in drug discovery and development settings. Adv Drug Deliv Rev. 46, 3–26.

Liu Z, Hamamichi S, Lee BD, Yang D, Ray A, Caldwell GA, Caldwell KA, Dawson TM, Smith WW, Dawson VL (2011). Inhibitors of LRRK2 kinase attenuate neurodegeneration and Parkinson-like phenotypes in Caenorhabditis elegans and Drosophila Parkinson’s disease models. Hum Mol Genet. 20, 3933–3942.

NCBI Resource Coordinators (2015). Database resources of the National Center for Biotechnology Information. Nucleic Acids Res. 43, D6–17.

Oh SW, Mukhopadhyay A, Svrzikapa N, Jiang F, Davis RJ, Tissenbaum HA (2005). JNK regulates lifespan in Caenorhabditis elegans by modulating nuclear translocation of forkhead transcription factor/DAF-16. Proc Natl Acad Sci U S A. 102, 4494–4499.

Pandey UB, Nichols CD (2011). Human disease models in Drosophila melanogaster and the role of the fly in therapeutic drug discovery. Pharmacol Rev. 63, 411–436.

Partridge FA, Tearle AW, Gravato-Nobre MJ, Schafer WR, Hodgkin J (2008). The C. elegans glycosyltransferase BUS-8 has two distinct and essential roles in epidermal morphogenesis. Dev Biol. 317, 549–559.

St Pierre SE, Ponting L, Stefancsik R, McQuilton P, FlyBase C (2014). FlyBase 102--advanced approaches to interrogating FlyBase. Nucleic Acids Res. 42, D780–788.

Tacutu R, Craig T, Budovsky A, Wuttke D, Lehmann G, Taranukha D, Costa J, Fraifeld VE, de Magalhães JP (2013). Human Ageing Genomic Resources: integrated databases and tools for the biology and genetics of ageing. Nucleic Acids Res. 41, D1027–D1033.

Trott O, Olson AJ (2010). AutoDock Vina: improving the speed and accuracy of docking with a new scoring function, efficient optimization, and multithreading. Journal of computational chemistry. 31, 455–461.

Tullet JM, Hertweck M, An JH, Baker J, Hwang JY, Liu S, Oliveira RP, Baumeister R, Blackwell TK (2008). Direct inhibition of the longevity-promoting factor SKN-1 by insulin-like signaling in C. elegans. Cell. 132, 1025–1038.

UniProt Consortium (2014). Activities at the Universal Protein Resource (UniProt). Nucleic Acids Res. 42, D191–198.

Wang R, Fang X, Lu Y, Wang S (2004). The PDBbind database: collection of binding affinities for protein-ligand complexes with known three-dimensional structures. Journal of medicinal chemistry. 47, 2977–2980.

Warner HR, Ingram D, Miller RA, Nadon NL, Richardson AG (2000). Program for testing biological interventions to promote healthy aging. Mech Ageing Dev. 115, 199–207.

Wilson MA, Rimando AM, Wolkow CA (2008). Methoxylation enhances stilbene bioactivity in Caenorhabditis elegans. BMC pharmacology. 8, 15.

Wurz RP, Pettus LH, Xu S, Henkle B, Sherman L, Plant M, Miner K, McBride H, Wong LM, Saris CJ, Lee MR, Chmait S, Mohr C, Hsieh F, Tasker AS (2009). Part 1: Structure-Activity Relationship (SAR) investigations of fused pyrazoles as potent, selective and orally available inhibitors of p38alpha mitogen-activated protein kinase. Bioorganic & medicinal chemistry letters. 19, 4724–4728.

Yang S, Long LH, Li D, Zhang JK, Jin S, Wang F, Chen JG (2015). beta-Guanidinopropionic acid extends the lifespan of Drosophila melanogaster via an AMP-activated protein kinase-dependent increase in autophagy. Aging Cell.

Ye X, Linton JM, Schork NJ, Buck LB, Petrascheck M (2014). A pharmacological network for lifespan extension in Caenorhabditis elegans. Aging Cell. 13, 206–215.

Zheng S-Q, Ding A-J, Li G-P, Wu G-S, Luo H-R (2013). Drug absorption efficiency in Caenorhbditis elegans delivered by different methods. PLoS One. 8, e56877–e56877.

